# Evaluation of ACMG Rules for In silico Evidence Strength Using An Independent Computational Tool Absent of Circularities on *ATM* and *CHEK2* Breast Cancer Cases and Controls

**DOI:** 10.1101/835264

**Authors:** Colin C. Young, Bing-Jian Feng, Colin B. Mackenzie, Elodie Girard, Donglei Hu, Yusuke Iwasaki, Yukihide Momozawa, Fabienne Lesueur, Elad Ziv, Susan L. Neuhausen, Sean V. Tavtigian

## Abstract

The American College of Medical Genetics and Genomics (ACMG) guidelines for sequence variant classification include two criteria, PP3 and BP4, for combining computational data with other evidence types contributing to sequence variant classification. PP3 and BP4 assert that computational modeling can provide “Supporting” evidence for or against pathogenicity within the ACMG framework. Here, leveraging a meta-analysis of *ATM* and *CHEK2* breast cancer case-control mutation screening data, we evaluate the strength of evidence determined from the relatively simple computational tool Align-GVGD. Importantly, application of Align-GVGD to these *ATM* and *CHEK2* data is free of logical circularities, hidden multiple testing, and use of other ACMG evidence types. For both genes, rare missense substitutions that are assigned the most severe Align-GVGD grade exceed a “Moderate pathogenic” evidence threshold when analyzed in a Bayesian framework; accordingly, we argue that the ACMG classification rules be updated for well-calibrated computational tools. Additionally, congruent with previous analyses of *ATM* and *CHEK2* case-control mutation screening data, we find that both genes have a considerable burden of pathogenic missense substitutions, and that severe *ATM* rare missense have increased odds ratios compared to truncating and splice junction variants, indicative of a potential dominant-negative effect for those missense substitutions.

## 1. Introduction

The American College of Medical Genetics and Genomics (ACMG) sequence variant classification guidelines include two criteria that enable integration of computational data with other evidence towards variant classification^1^. These criteria, known as “PP3” and “BP4”, assert that computational modeling can provide supporting evidence for or against pathogenicity, respectively. There is a thought in the field, however, that computational modeling should be able to provide stronger than “supporting” evidence for or against pathogenicity, but rigorous validation of computational tools is difficult to perform due to hidden multiple testing, circularities between training sets used to create/train/calibrate the computational tool, and hidden use of other ACMG evidence categories.

To test the hypothesis that a computational tool can be used to generate stronger evidence for or against pathogenicity, we wanted to use a tool that is free from the problems described above. Align-GVGD is a relatively simple computational tool that assigns missense severity based on the physicochemical difference between a missense amino acid and the range of variation observed at its position in a suitably informative protein multiple sequence alignment (PMSA)^2–4^. The only inputs to Align-GVGD are a table of amino acid sidechain composition, polarity, and volume data compiled in 1974^2^, a list of missense substitutions of interest, and a user-supplied PMSA. The program does not call any external data and thus does not access any ACMG evidence categories other than PP3 and BP4.

To validate using Align-GVGD, we performed a meta-analysis of breast cancer case/control mutation screening data from two genes, *ATM* and *CHEK2*. These two genes are securely classified as intermediate-risk susceptibility genes, and it is clear that at least some missense substitutions in both *ATM* and *CHEK2* confer increased risk for breast cancer^5–14^. In this study, we combine three recent studies that included mutation screening of both *ATM* and *CHEK2* in cases and controls: Girard et al. (2019), Weitzel et al. (2019), and Momozawa et al. (2018)^15–17^. Combined, these studies include 9,311 breast cancer cases and 13,629 controls. The user-supplied PMSA that we used to score the observed rare missense substitutions (rMS) were created for and then left unchanged since our 2009 and 2011 studies of *ATM* and *CHEK2*, respectively^7,11^. Additionally, the calibration of Align-GVGD to create four graded analysis categories of increasing predicted pathogenicity was performed in 2008 using data from *BRCA1* and *BRCA2* mutation screening studies^4,18^. Consequently, there are no circularities between the creation, calibration, and required data inputs to Align-GVGD or the *ATM* and *CHEK2* data evaluated here.

## 2. Methods

### Study Characteristics for Meta-Analysis

The studies chosen for this work were selected after a literature search in June 2019 and were selected to meet three criteria: 1) they were bona fide breast cancer case-control studies; 2) all of the coding exons were sequenced; and 3) contained more than 1,000 cases and 1,000 controls. We chose to focus on cancer case/control mutation screening studies that included screening for rMS in both *ATM* and *CHEK2* (summarized in Table 1). The first study published by Girard et al. (2019) includes women who presented with breast cancer, had a sister affected with breast cancer, and were screened and selected based on the absence of *BRCA1/2* pathogenic variants. The second study was a case/control mutation screening performed in self-identified Hispanic women published by Wietzel et al. (2019). Participants in this study were also screened against having a *BRCA1/2* pathogenic variant. The third study was a large scale study performed on Japanese women from Biobank Japan published by Momozawa et al. (2018). In the Momozawa et al. data, one case subject was excluded for both genes as the subjects also had a pathogenic rMS variant in *BRCA2*, meaning the number of cases analyzed in this work for both genes was 7,050.

**Table 1.** Case/Control Mutation Screening Study Characteristics

| Study | Cases | Controls | ATM<br>Carrier<br>Subjects | CHEK2<br>Carrier<br>Subjects | Distinct ATM<br>Variants | Distinct CHEK2<br>Variants |
| --- | --- | --- | --- | --- | --- | --- |
| Girard et al. | 1207 | 1199 | 221 | 94 | 126 | 50 |
| Weitzel et al. | 1054 | 1189 | 232 | 63 | 112 | 34 |
| Momozawa et al. | 7050<sup>a</sup> | 11241 | 1280 | 406 | 227 | 63 |
| Total | 9311 | 13629 | 1733 | 563 | 368<sup>b</sup> | 110<sup>b</sup> |

<sup>a</sup> One case was excluded from the original study for each gene due to the subject also carrying a pathogenic BRCA2 mutation
<sup>b</sup> Some variants were observed in multiple studies

In collaboration with each study’s authors, we identified any subjects who were observed having have two different rare variants of interest – either protein truncating variants or rMS – in the same gene. In these cases, the subject was categorized based on the most severe rMS variant. We also note that some carriers of in-frame deletions, of deletions in regions that demonstrated high variability in the PMSA, or of frameshifts occurring in the final few exons of either gene and not expected to cause nonsense mediated mRNA decay, were classified as being missense carriers or excluded as non-carriers depending on the predicted severity. Detailed information on how these variants were assigned can be found in the supplemental data.

### Align-GVGD

The hand-curated PMSAs used for this analysis are the same alignments used in our *ATM* and *CHEK2* case/control mutation studies published earlier (available at http://agvgd.hci.utah.edu/alignments.php); we are specifically using the same alignments to minimize hidden multiple testing. Each rMS variant was compared against the PMSA from human to sea urchin to predict severity. Align-GVGD assigns rMS to a series of seven grades of increasing predicted severity; C0, C15, C25, C35, C45, C55, and C65. For most analyses, these Align-GVGD grades were collapsed into four categories (C0; C15,25; C35-55; and C65) as in our previous *BRCA1/2* analysis^4^. To maintain compatibility with our PMSA, *CHEK2* variants were annotated using transcript form NM_007194.3; transcript form NM_000051.3 was used for *ATM*.

### Allele Frequency Analysis

rMS were also stratified by allele frequency as measured in the non-cancer population database available from GnomAD. The Hereditary Breast and Ovarian Cancer Variant Classification Expert Panel (HBOC VCEP) has preliminarily proposed that the ACMG allele frequency-based stand-alone (BA1) and strong benign (BS1) evidence for *ATM* and *CHEK2* could be: BA1, above an allele frequency of 0.005; and BS1, above an allele frequency 0.0005, but less than 0.005 [Marcy Richardson and Amanda Spurdle, personal communication]. Our analysis also requires a BS1_moderate threshold, for which we have adopted the HBOC VCEP *BRCA2* BS1 threshold of ≤0.00014. The resulting frequency strata are summarized in Table 2.

**Table 2.** Allele Frequency Evidence Thresholds in Population Genetic Databases as Set by HBOP VCEP

| Threshold Criteria | BRCA1/2 | ATM/CHEK2 |
| --- | --- | --- |
| Stand-alone Benign (BA1) | 0.0014 | 0.005 |
| Strong Benign (BS1) | 0.00014 | 0.0005 |
| Benign Moderate | >2 observations | 0.00014<sup>a</sup> |
| Indeterminate | 1-2 observations | - |
| PM2 Supporting | Absent | - |

<sup>a</sup> Adopted for this work

*ATM* and *CHEK2* rMS were cross-referenced with the GnomAD database to determine the continental-level allele frequencies; African, Latino, Non-Finnish European, East Asian, and South Asian allele frequencies were used. rMS absent from GnomAD were assigned an allele frequency of zero. Probably because the number of Japanese cases and controls in the third study was similar to the total number of East Asians, and much greater than the number of Japanese alleles in GnomAD, allele frequencies for East Asians from GnomAD were not representative of our data. To account for this, we added all the control alleles and a proportion of the case allele counts such that the cases accounted for 10% of the Japanese subjects added from Study 3 to the East Asian sequence variant data recorded in GnomAD. We then recalculated the East Asian allele frequencies based on this higher number of East Asian subjects and observed sequence variants.

The highest allele frequency for any continental level race/ethnicity was then used as the maximum frequency for each variant. The allele frequencies were sorted into three different frequency bins: the first frequency bin contained variants with an allele frequency between 0 and 0.00014, the second frequency bin contained variants with an allele frequency between 0.00014 and 0.0005, the third frequency bin contained variants between 0.0005 and 0.005. Subjects only carrying a variant with an allele frequency above 0.005 were included in this study as a noncarrier.

### Statistical Methods

A database for each gene was constructed with an entry for each individual subject. The database indicated whether a subject had a truncating or splice junction variant (T+SJV), rMS variant, case/control status, annotation for study, Align-GVGD grade (0 for non-carriers, 1-7 for C0 up to C65), Align-GVGD category, allele frequencies for each variant in African, Latino, East Asian, non-Finnish European, and South Asian subjects, and a designation indicating the highest continental allele frequency bin assigned for each subject. For the purposes of our analysis, study was used as categorical variable for all logistic regressions, and due to the specific nature the of ethnicities in each study, should account for differences in ethnicity.

Logistic regressions on the subject populations for each gene were performed on subjects carrying T+SJV against noncarriers, without factoring in allele frequency, to determine the odds ratio (OR) for T+SJV carriers. Similarly, logistic regressions were also performed on rMS carriers in each frequency bin for each individual Align-GVGD category (*e*.*g*., C0 or C35-55). Logistic regression trend tests were performed on the rMS carriers of the seven Align-GVGD grades where the Align-GVGD grade was 0-7 (0 for noncarriers, 1 for C0 carriers, 2 for C15 carriers etc.). The rMS in each of the four Align-GVGD categories and in each frequency bin were then analyzed individually, excluding T+SJV carriers. For these analyses, noncarriers were assigned as a “0,” while subjects in each of four Align-GVGD grade categories were assigned as “1” for being rMS carriers. Each Align-GVGD category was analyzed separately, excluding the subjects in the other categories and frequency bins. All logistic regressions were performed using the logit function in the Python StatsModels toolbox.

Odds in favor of pathogenicity for the four Align-GVGD missense substitution categories were estimated through a three-step procedure^19^. First, separately for *ATM* and *CHEK2*, we estimated an OR for each of the four Align-GVGD categories by taking a weighted average of the ORs obtained for each of the four categories in each of the three allele frequency strata. Second, we estimated a maximum likelihood proportion of pathogenic variants for the four Align-GVGD categories. To do this, we used the total number of missense substitutions placed in each category along with the categorical OR for that category, the OR for T+SJV variants and a theoretical OR of 1.00 for a “pure” set of benign variants. We then determined the ratio of variants with the OR of the T+SJV bin to variants with OR = 1.00 that best approximated the observed categorical OR for each category. In the instances where the OR of an rMS category was higher than the OR for T+SJVs, this proportion was set to n/(n+1), where n is the number of rMS in the bin (thus allowing the proportion to approach but not exceed 1.0). Finally, we estimated the odds in favor of pathogenicity for the four Align-GVGD categories. To do this, we treated the estimated proportion of pathogenic variants in the overall set of rMS as a prior probability (*P_1_*) and the proportion estimated for each of the four Align-GVGD categories as a posterior probability (*P_2_*). Odds path were then estimated as:

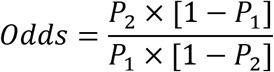

## 3. Results

### Analysis of Truncating and Splice Junction Variants

As in our previous analyses on *ATM* and *CHEK2*, variants that introduce significant frameshifts, premature stop codons, and those expected to destroy a splice junction and cause nonsense mediated decay were pooled and analyzed for each gene individually. Allele frequency of T+SJVs was not included as a factor in this analysis, due to the high continental allele frequency of some known founder pathogenic variants (e.g., c.1100delC in *CHEK2*) and also because the ACMG PVS1 criterion (null variant) outweighs the BS1 criterion (allele frequency greater than expected for disorder).

For *ATM* there was a total of 72 T+SJV, with 42 variants appearing in cases and 28 variants appearing in controls. For *CHEK2* there was 55 T+SJV, 39 in cases and 16 in controls. Logistic regression on the presence of T+SJVs against noncarriers returned an OR=2.01 (P=0.0046) for *ATM* variants and OR=3.19 (P=0.00010) for *CHEK2* variants. These data are displayed in Table 3.

**Table 3.**
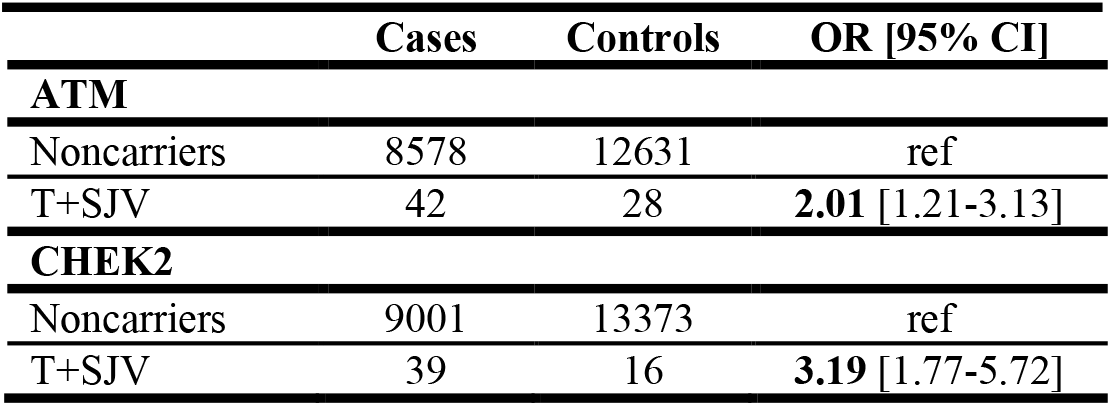
Analysis of Truncating and Splice Junction Variants

### Analysis of rare Missense Substitutions

There are a number of computational tools that are commonly used to group rMS variants by predicting severity of the variant. For our analyses, Align-GVGD, along with previously published PMSAs, were used to pool the rMS variants of both genes into four categories for analysis. Of note, these are the same categories previously used to determine sequence analysis-based prior probabilities of pathogenicity for key domain rMS in *BRCA1* and *BRCA2*^4^.

Progressing from the lowest allele frequency bin to the higher frequency bins, the number of distinct rMS per bin decreased. In general, even as the number of distinct rMS decreased, the number of subjects in the higher frequency bins increased because more common variants were found in more subjects. For *CHEK2*, the number of rMS carrying subjects in each frequency bin increased from 156 in frequency bin 1 to 179 in frequency bin 2, and to 231 in frequency bin 3. For *ATM* the counts were 464, 273, and 985 subjects in rMS frequency bins 1, 2, and 3, respectively. A summary of the ORs for each gene and in each frequency bin is summarized in Table 4.

**Table 4.** Summary of Odds Ratios Measured in Each Align-GVGD Category and Frequency Bin

| Align-GVGD Category | ATM<br>Distinct rMS<sub>a</sub> | ATM<br>OR [95% CI] | CHEK2<br>Distinct rMS<sub>a</sub> | CHEK2<br>OR [95% CI] |
| --- | --- | --- | --- | --- |
| **Frequency Bin 1 ($0 \leq \text{Allele Frequency} < 0.00014$)** |  |  |  |  |
| C0 | 182 | **1.30** [1.02-1.66] | 37 | 1.17 [0.70-1.96] |
| C15,25 | 37 | **1.80** [1.07-3.02] | 15 | 2.28 [0.95-5.46] |
| C35-55 | 28 | 1.76 [0.93-3.32] | 8 | **4.08** [1.29-12.85] |
| C65 | 27 | **4.41** [1.89-10.28] | 15 | **4.32** [1.43-13.01] |
| Any rMS | 274 | **1.52** [1.24-1.86] | 75 | **1.87** [1.29-2.72] |
| **Frequency Bin 2 ($0.00014 < \text{Allele Frequency} < 0.0005$)** |  |  |  |  |
| C0 | 36 | 0.89 [0.68-1.17] | 9 | **1.59** [1.03-2.31] |
| C15,25 | 2 | 1.64 [0.27-19.84] | 5 | **2.52** [1.48-4.29] |
| C35-55 | 4 | 0.96 [0.41-2.27] | 1 | **3.46** [1.21-10.00] |
| C65 | 6 | 1.04 [0.56-1.93] | 3 | 0.31 [0.07-1.43] |
| Any rMS | 48 | 0.91 [0.71-1.17] | 18 | **1.82** [1.34-2.44] |
| **Frequency Bin 3 ($0.0005 < \text{Allele Frequency} < 0.005$)** |  |  |  |  |
| C0 | 28 | 0.88 [0.76-1.03] | 11 | **1.41** [1.01-1.96] |
| C15,25 | 8 | 0.90 [0.66-1.23] | 3 | 1.03 [0.60-1.74] |
| C35-55 | 2 | 0.54 [0.13-2.15] | 3 | **5.40** [1.55-18.82] |
| C65 | 4 | 1.11 [0.73-1.68] | 0 | – |
| Any rMS | 46 | 0.90 [0.79-1.03] | 17 | **1.42** [1.08-1.85] |

<sup>a</sup> The number of noncarriers was the same for each gene as those used for the T+SJV's in Table 3.

For *ATM*, there were 274 distinct rMS variants in the lowest frequency bin; overall, these were associated with OR=1.52 (P=0.000042). Stratifying into the four Align-GVGD categories, there were 182 rMS in the C0 category, 37 in the C15,25 category, 28 in the C35-55 category, and 27 in the C65 category. Across these categories, ORs increased from 1.30 (P=0.033) for C0 rMS, 1.80 (P=0.026) for C15,25 rMS, 1.76 (P=0.083) for C35-55 rMS, and 4.41 (P=0.00059) for C65 rMS. Additionally, a trend test across the seven Align-GVGD grades, which is against the null hypothesis of no change in OR with increasing grade of rMS, yielded a ln(OR) increase of 0.18 per grade (P_trend_=0.000003).

Overall, the 75 *CHEK2* rMS in the lowest frequency bin were associated with OR=1.87 (P=0.0010). Stratifying in the Align-GVGD categories, there were 37 distinct rMS in the C0 category, 15 variants in the C15,25 category, 8 variants in the C35-55 category, and 15 variants in the C65 category. C0 rMS variants had an OR=1.17 (P=0.54), C15,25 variants had an OR=2.28 (P=0.064), C35-55 variants had an OR=4.08 (P=0.016), ending at OR=4.32 (P=0.010) for C65 missense substitutions. The trend test across the seven Align-GVGD grades was also significant (ln(OR)=0.24 per grade, P_trend_=0.000088).

rMS with allele frequencies falling into the 2nd and 3rd frequency bins were assessed similarly (Table 4). For *ATM*, there appeared to be a strong inverse relationship between allele frequency and evidence of risk for breast cancer; indeed, all of the individually significant Align-GVGD results were restricted to frequency bin 1. In contrast, for *CHEK2*, there were multiple individually significant results in each of the three allele frequency bins, and the overall ORs for rMS in the three frequency bins were all significantly elevated. Even so, one *CHEK2* C65 rMS in frequency bin 2, p.P213H, was observed in 0 cases and 7 controls in Momozawa et al., which resulted in an OR for C65 rMS in bin 2 of 0.31 (P=0.136).

Our previous work analyzing rMS variants for *ATM* demonstrated an increased risk of breast cancer for rMS and in-frame indels falling in the kinase-active carboxy-third of the protein (after Ile1960) as opposed to variants falling into the first two thirds of the protein (before Ile1960). Here, we did not observe a significant difference between the ORs for rMS variants in these two intervals (data not shown).

### From Frequentist Odds Ratios to Bayesian Odds Path

The ACMG sequence variant classification guidelines employ a series of strength of evidence categories; recently, we fit these to a Bayesian framework and in so doing worked out thresholds for each category expressed as odds in favor of pathogenicity (Odds Path)^20^. Accordingly, we estimated Odds Path for the Align-GVGD categories as applied to *ATM* and *CHEK2*; these are summarized in Table 5.

**Table 5.** Align-GVGD Category Odds Ratios Converted to Odds of Pathogenicity

| AGVGD Category | ATM<br>Odds of<br>Pathogenicity | CHEK2<br>Odds of<br>Pathogenicity | Combined<br>Odds of<br>Pathogenicity | Evidence<br>Strength | Evidence<br>Threshold |
| --- | --- | --- | --- | --- | --- |
| C0 | 0.41 | 0.25 | 0.38 | Supporting<br>Benign | Odds<0.38 |
| C15,25 | 3.23 | 1.99 | 2.82 | Supporting<br>Pathogenic | 2.08<Odds<4.3 |
| C35-55 | 2.62 | 21.85 | 7.63 | Moderate<br>Pathogenic | 4.33<Odds<18.7 |
| C65 | 67.74 | 32.77 | 56.29 | Strong<br>Pathogenic | Odds>18.7 |

For *ATM* variants Odds Path were estimated to be 0.41 for C0 rMS, 3.23 for C15,25 rMS, 2.62 for C35-55 rMS, and 67.74 for C65 rMS. For *CHEK2* Odds Path were estimated to be 0.25 for C0 rMS, 1.99 for C15,25 rMS, 21.85 for C34-55 rMS and 32.77 for C65 rMS. Using weighted averages to combine across the two genes, Odds Path were estimated to be 0.38 for C0 rMS, 2.82 for C15,25 rMS, 7.63 for C35-55 rMS, and 56.29 for C65 rMS.

The combined gene analysis shows a clear trend of increasing evidence strength as a function of Align-GVGD category. Odds Path for C0 rMS fall into the ACMG “Supporting Benign” category; C15,25 rMS are “Supporting Pathogenic”; C35-55 rMS are “Moderate Pathogenic”, and the C65 rMS reach the “Strong Pathogenic” category.

## 4. Discussion

For this analysis, our primary objective was to test the hypothesis that a computational tool that is free of ACMG evidence types other than PP3 and BP4 can produce stronger than “Supporting” evidence of pathogenicity. Align-GVGD relies only on the physicochemical deviations of amino acid substitutions from the range of variation observed at their position in a curated PMSA, and therefore meets this criterion. As applied to *ATM* and *CHEK2*, the calibration of Align-GVGD has no hidden multiple testing or circularities, making it an ideal candidate to evaluate the strength of evidence provided by a computational tool.

The results of the odds of pathogenicity calculations demonstrate that a specific computational tool can provide stronger than supporting evidence of pathogenic effect. In a weighted average across *ATM* and *CHEK2* and across the three bins with allele frequency below the PVS1 threshold of 0.005, both the Align-GVGD C35-C55 and C65 categories produced Odds Path that were above the ACMG “Moderate Pathogenic” threshold of Odds Path=4.33. At the same time, the Align-GVGD C0 category produced Odds Path that fall in the ACMG “Supporting Benign” range; these results are similar to those that we recently reported for TP53^21^, though generated within a cleaner analytic framework. Based on the analysis here, we argue that PP3 should be updated to allow a “PP3_Moderate” strength of evidence for well-calibrated gene-computational tool combinations.

While the main focus of this work was to examine strength of evidence attributable to the ACMG PP3 code, the case-control data analyzed here also allowed us to reevaluate findings from our previous *ATM* and *CHEK2* mutation screening studies, which had been performed with independent series of case and control subjects. Overall, we find that the number of cases with rMS severity grades of C35-55 and C65 across frequency bins to be on par with the number of cases carrying T+SJVs, and have similar or elevated ORs to those of T+SJVs. This echoes our previous result and underlines the need for accelerated classification of rMS in these genes.

For *ATM*, we previously reported that: (1) most evidence for pathogenic rMS was confined to the FAT, kinase, and FATC domains, roughly from amino acid 1960 until the end of the protein, and (2) that C65 missense substitutions in ATM confer greater risk of breast cancer than do T+SJVs. The first point was not supported by our current results – in these data, there is no evidence for a difference in OR for rMS falling before (OR=1.64) or after amino acid 1960 (OR=1.66). On the other hand, the OR for C65 missense substitutions in frequency bin 1 was more than twice the OR of T+SJVs (4.41 vs 2.01, respectively). Although the result is not statistically significant, it remains noteworthy because the effect is distributed across many rMS rather than being largely due to the single known pathogenic p.V2424G founder mutation^13^ (which was carried by only one subject – a control – in this study). This relatively high OR could indicate a dominate-negative effect, which warrants mechanistic studies.

The ACMG allele-frequency related evidence codes BA1 and BS1 imply a strong inverse relationship between allele frequency and risk. That inverse relationship is clearly evident in the *ATM* data, where essentially all risk associated with rMS is limited to those in frequency bin 1. In contrast, for *CHEK2*, there is evidence for pathogenic rMS in frequency bins 2 and 3. We speculate that this difference may trace to the fact that *ATM* is a recessive susceptibility gene for an early onset, high-mortality disease that limits reproductive fitness for homozygotes and compound heterozygotes, whereas *CHEK2* is not. However, we also note that there is evidence that some *CHEK2* alleles that are mildly pathogenic for breast cancer confer resistance to smoking-related cancers^22,23^; if some dysfunctional *CHEK2* sequence variants are involved in balancing selection, then the expected pattern of allele frequency vs risk of breast cancer could be distorted.

## 5. Conclusion

Here we demonstrate that a computational tool that is absent of other ACMG evidence types, circularities, and hidden multiple testing can be used to predict stronger than PP3 “Supporting_Pathogenic” evidence for the breast cancer susceptibility genes *ATM* and *CHEK2*. The point is not that the specific computational tool used here, Align-GVGD, is superior to others; rather, the forward looking question is whether the strength of PP3 is only under-estimated for a few genes versus under-estimated in general. We argue that PP3 could be updated to include the possibility of moderate strength of evidence for well-calibrated gene-computational tool combinations, provided that the computational tool does not leverage other evidence codes and the calibration is free of circularities.

Additionally, we were able to reevaluate some of our findings from previous analyses of *ATM* and *CHEK2* variants. In particular, we find (1) statistical evidence that *ATM* and *CHEK2* both have a considerable burden of pathogenic missense substitutions, and (2) Align-GVGD C65 missense substitutions in *ATM* continue to show evidence of increased ORs as compared to T+SJVs, indicating a potential dominate-negative effect.

